# Tximeta: reference sequence checksums for provenance identification in RNA-seq

**DOI:** 10.1101/777888

**Authors:** Michael I. Love, Charlotte Soneson, Peter F. Hickey, Lisa K. Johnson, N. Tessa Pierce, Lori Shepherd, Martin Morgan, Rob Patro

## Abstract

Correct annotation metadata is critical for reproducible and accurate RNA-seq analysis. When files are shared publicly or among collaborators with incorrect or missing annotation metadata, it becomes difficult or impossible to reproduce bioinformatic analyses from raw data. It also makes it more difficult to locate the transcriptomic features, such as transcripts or genes, in their proper genomic context, which is necessary for overlapping expression data with other datasets. We provide a solution in the form of an R/Bioconductor package tximeta that performs numerous annotation and metadata gathering tasks automatically on behalf of users during the import of transcript quantification files. The correct reference transcriptome is identified via a hashed checksum stored in the quantification output, and key transcript databases are downloaded and cached locally. The computational paradigm of automatically adding annotation metadata based on reference sequence checksums can greatly facilitate genomic workflows, by helping to reduce overhead during bioinformatic analyses, preventing costly bioinformatic mistakes, and promoting computational reproducibility. The tximeta package is available at https://bioconductor.org/packages/tximeta.

## Introduction

An RNA-seq data analysis often involves quantification of sequence read data with respect to a set of known reference transcripts. These reference transcripts may be downloaded from a database such as GENCODE, Ensembl, or RefSeq [1–3] in the form of nucleotide sequences in FASTA format and/or transcript locations in a genome in GTF/GFF (gene transfer format / general feature format). Alternatively a novel set of reference transcripts may be derived as part of the data analysis. The provenance of the reference transcripts, including their source and release number, can be considered critical metadata with respect to the processed data, that is, the files with quantification information. Without information about the reference provenance, computational reproducibility — re-performing the analysis with the same data and code and obtaining the same result [4] — may be difficult or impossible. Reproducibility has been set as a high-level goal for all NIH-funded research [5, 6], and so as developers of bioinformatic tools, we should design software that promotes and facilitates computational reproducibility. Manually keeping track of critical pieces of metadata throughout a long-term bioinformatic project is tedious and error prone; still, manual metadata tracking is a common practice in RNA-seq bioinformatics. For example, a common approach to tracking the reference transcripts that were used during quantification would be to keep a `README` file in the same directory as the quantification data, with information about the provenance of the reference transcripts.

In addition to impeding computational reproducibility, missing or wrong metadata can potentially lead to serious errors in downstream analysis: if quantification data are shared with genomic coordinates but without critical metadata about the genome version, computation of overlaps with other genomic data with mis-matching genome versions can lead to faulty inference of overlap enrichment. Additional annotation tasks, such as conversion of transcript or gene identifiers, or summarization of transcript-level data to the gene level is made more difficult when the reference provenance is not known. Kanduri *et al.* [7] documented issues surrounding the lack of provenance metadata for BED, WIG, and GFF files, and described this problem as a “major time thief” in bioinformatics. Likewise, Simoneau and Scott [8] described information on genome assembly and annotation as “essential” for describing the computational analysis of RNA-seq data, and contended that, “no study using RNA-seq should be published without these methodological details.”

A number of frameworks have been proposed that would solve the problem of tracking provenance in a bioinformatic analysis – provenance in the narrow sense defined above, encompassing the source and release information of the reference sequence – as well as in a larger sense of tracking the state of all files, including data and metadata and any software used to process these files, throughout every step of an analysis. We will first review frameworks for tracking provenance of reference sequences, and secondly describe more general frameworks. The CRAM format, developed at the European Bioinformatics Institute, involves computing differences between biological sequences and a given reference so that the sequences themselves do not need to be stored in full within an alignment file [9]. Because the specific reference used for compression is critical for data integrity, CRAM includes checksums of the reference sequences as part of the file header. A partner utility called refget has been developed in order to allow for programmatic retrieval of the reference sequence from a computed checksum, which acts as an identifier of the reference sequence [10]. A similar approach is taken by the Global Alliance for Genomics and Health’s (GA4GH) Variation Representation Specification (VR-Spec) [11], which uses a hashed checksum (or “digest”) to uniquely refer to molecular variation, and by the seqrepo python package for writing and reading collections of biological sequences [12]. The NCBI Assembly database takes a different approach, by assigning unambiguous identifier strings (though not computed via a hash function) to sets of sequences comprising specific releases of a genome assembly [13]. Knowing the identifier is therefore sufficient to know the full set of sequences in the assembly. Another approach to reduce manual metadata tracking associated with a number of reference sequences is Refgenie. Refgenie is a tool that helps with management of bundles of files associated with reference genomes, and facilitates sharing provenance information across research groups, in that the generation of resources is scripted [14]. Arkas, ARMOR, pepkit, and basejump are all frameworks for automating bioinformatic analyses, where reference provenance is specified in configuration files and correct metadata can therefore be assembled and attached programmatically to downstream outputs [15–18].

In 2015, Belhajjame *et al.* [19] introduced the concept of a “Research Object”, an aggregation of data and supporting metadata produced within a specified scientific workflow. Their formulation was system-neutral, describing the requirements for production of a Research Object. The requirements touch on topics introduced above, such as the need to preserve data inputs, software versions, as well as traces of the provenance of data as it moves through the scientific workflow. Belhajjame *et al.* [19] summarized literature in the field of computational reproducibility and efforts toward extensive provenance tracking. The developers of the Common Workflow Language (CWL) [20] have defined a profile, CWLProv, for recording provenance through a workflow run, and have a number of implementations, including within cwltool [21]. The developers of CWLProv emphasized the importance of tracking versions of input data, such as reference genomes or variant databases in a scientific workflow, and they suggested to use and store stable identifiers of all data and software, as well as the workflow itself. As identifiers play such a crucial role in assuring reproducibility of workflows, the developers of CWLProv recommended the use of hashed checksums for identifiers of data (including any reference sequence), similar to the use of checksums in the CRAM format and VR-Spec, for identifying the reference or variant sequences. Gruning *et al.* [22] recommended combining systems such as Galaxy for encapsulating analysis tools with systems for tracking and capturing parameters and source data provenance to provide full computational reproducibility.

Here we describe an R/Bioconductor package, tximeta, for identification of reference transcript provenance in RNA-seq analyses via sequence checksums. It is situated among other solutions for facilitating computational reproducibility described above, with some automation of routine tasks, such as conversion of transcript and gene names, but short of full automation of analyses as in Arkas and ARMOR. Tximeta captures the versions of the software packages used in import of quantification data, but does not provide full provenance tracking throughout downstream tasks as in the Research Object specification or in CWLProv. One desirable aspect of tximeta is that – through the use of hashed checksums of reference transcripts and lookup operations similar to those performed by refget – our implementation can be used to identify the reference provenance *post hoc* on various shared or public datasets, regardless of whether the original analyst kept or shared accurate records of the reference transcripts that were used. Therefore it can provide some utility for bioinformatic analysts without requiring full buy-in of a particular workflow execution framework. Tximeta is similar in implementation to the CRAM format in the use of hashed checksums, but identifies the transcript sequences used during sample quantification rather than the genome sequence used during alignment. We see tximeta as a piece of a larger effort to create software systems that are “more amenable to reproducibility” [23].

## Design and Implementation

Tximeta has been developed to work with output from Salmon or Alevin quantification tools [24, 25], although the implementation could be extended to other quantification tools that store the appropriate hashed checksum within the index and propagate this checksum to the sample output metadata. Without loss of generality, we describe the implementation referring to Salmon quantification data below. A diagram of the following workflow is shown in Figure 1.

**Fig 1.**
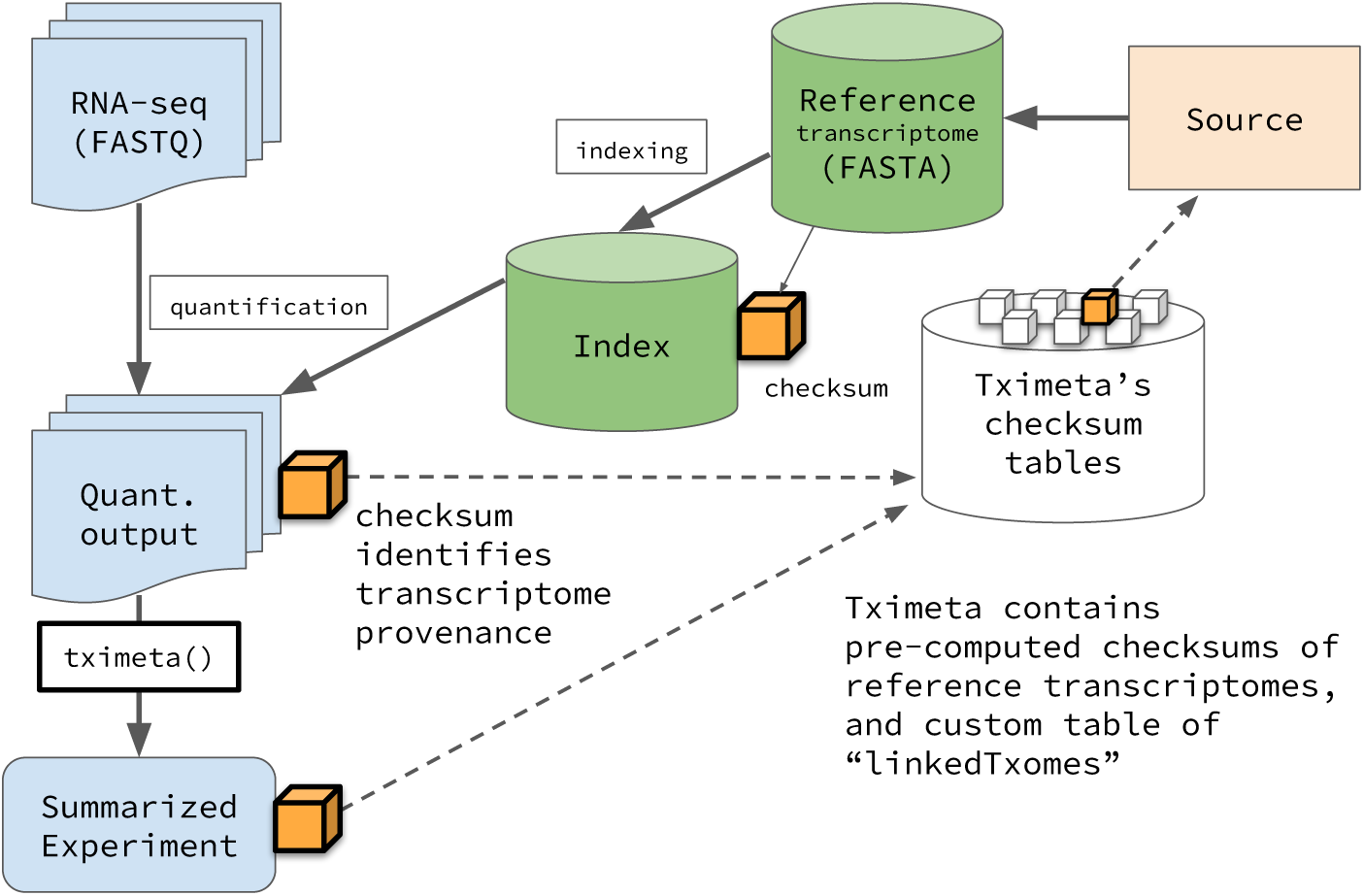
Flowchart of Salmon quantification followed by tximeta. The quantification and import pipeline results in a SummarizedExperiment object with reference transcript provenance metadata added by tximeta (see Design and Implementation).

During the indexing step, Salmon computes the hashed checksum of the cDNA sequence of the reference transcripts. The set of reference transcripts provided to Salmon will be referred to in this text as the *transcriptome*, although we note that the reference is not necessarily equal to the complete set of possible RNA transcripts in the sample. Currently, both the SHA-256 and SHA-512 [26] checksums are computed on the reference cDNA sequences alone, with transcript sequences concatenated together with the empty string (the SHA-256 checksum is currently taken as the primary identifier). Future implementations of Salmon and tximeta may use alternate hash functions for compatibility with larger efforts toward stable identifiers for sequence collections, for example, computing a hashed checksum over a lexicographically sorted set of checksums for each transcript cDNA sequence, which would provide order-invariance for the collection identifier. During quantification of a single sample, Salmon embeds the transcriptome index checksum in a metadata file associated with the sample output. For each sample, Salmon outputs a directory with a specific file structure, including files with quantification information as well as others with important metadata about the parameters. The entire directory, not just the text file with the quantification information, should be considered the output of the quantification tool.

During import of quantification data into R/Bioconductor [27], leveraging the existing tximport package [28], tximeta reads the quantification data, as well as the transcriptome index checksum, and compares this checksum to a hash table of pre-computed checksums of a subset of commonly used reference transcriptomes (human, mouse, and fruit fly reference transcripts from GENCODE, Ensembl, and RefSeq, see Table 1), as well as to a custom hash table which will be described below. Tximeta verifies that the checksum and therefore the reference transcriptome sequence is identical across all samples being imported. If there is a match of the checksum among the pre-computed checksums or in the custom hash table, tximeta will begin to compile additional relevant metadata. Depending on whether the checksum has been seen by tximeta before, one of two steps will occur:

**Table 1.** Pre-computed reference transcripts checksums in tximeta as of Fall 2019.

| Source | Organism | Releases | Transcript sequence file |
| --- | --- | --- | --- |
| GENCODE | *Homo sapiens* | 23 – 31 | transcripts.fa |
| GENCODE | *Mus musculus* | M6 – M22 | transcripts.fa |
| Ensembl | *Homo sapiens* | 76 – 97 | *.cdna.all.fa (NR) |
| Ensembl | *Mus musculus* | 76 – 97 | *.cdna.all.fa (NR) |
| Ensembl | *Drosophila melanogaster* | 79 – 97 | *.cdna.all.fa (NR) |
| Ensembl | *Homo sapiens* | 76 – 97 | *.cdna.all.fa + *.ncrna.fa |
| Ensembl | *Mus musculus* | 76 – 97 | *.cdna.all.fa + *.ncrna.fa |
| Ensembl | *Drosophila melanogaster* | 79 – 97 | *.cdna.all.fa + *.ncrna.fa |
| RefSeq | *Homo sapiens* | p1 – p12† | *_rna.fa |
| RefSeq | *Mus musculus* | p2 – p5† | *_rna.fa |

The set of pre-computed checksums span the stable releases from these sources for the years 2015 – 2019. (NR) - not recommended: we recommend combination of coding and non-coding transcripts for accurate RNA-seq quantification; † - RefSeq assembly versions p13 and p6 for human and mouse respectively are currently “latest”, and are subject to sequence updates under the same assembly version, and so not stable releases.

- (First time) - Tximeta attempts to download the appropriate GTF/GFF file via FTP and parse it using Bioconductor packages. GENCODE and RefSeq GTF/GFF files are parsed by GenomicFeatures [29], while Ensembl GTF files are parsed by ensembldb [30]. Tximeta then creates a locally cached SQLite database of the parsed GTF/GFF file, as well as a GRanges object of the transcript locations [29]. The local cache is managed by the BiocFileCache Bioconductor package [31].
- (Subsequently) - Tximeta loads the locally cached versions of metadata (the transcript ranges, or additionally the SQLite database on demand for further annotation tasks).

After loading the appropriate annotation metadata, tximeta outputs a SummarizedExperiment object [29], a class in the Bioconductor ecosystem which stores multiple similarly shaped matrices of data, or “assays”, including the estimated read counts, effective transcript lengths, and estimates of abundance (in transcripts per million, TPM). By convention, rows correspond to genomic features (e.g. transcripts or genes), while columns correspond to samples. In addition, the rows of the matrices are linked to transcript ranges, embedded in an appropriate genome version (e.g. GRCh38) including chromosome names and lengths. If tximeta did not find a matching transcriptome in the hash table then a non-ranged SummarizedExperiment will be returned as the function’s output, as the location and context of the transcript ranges are not known to tximeta. Comparison of ranges across genome versions, or without properly matching chromosomes, will produce an error, leveraging default functionality from the underlying GenomicRanges package [29]. Metadata about the samples, if provided by the user, is automatically attached to the columns of the SummarizedExperiment object. Additional metadata attached by tximeta includes all of the per-sample metadata saved from Salmon (e.g. library type, percent reads mapping, etc.), information about the reference transcriptome and file paths or FTP URLs for the source file(s) for FASTA and GTF/GFF, and the package versions for tximeta and other Bioconductor packages used during the parsing of the GTF/GFF. At any later point in time, annotation tasks can be performed by on-demand retrieval of the cached databases, for example summarization of transcript-level information to the gene level, conversion of transcript or gene identifiers, or addition of exon ranges.

A key aspect of the tximeta workflow described here is that it does not rely on self-reporting of the reference provenance for *post hoc* identification of the correct metadata. An exception to this rule is the case of a *de novo* constructed transcriptome, or in general, use of a transcriptome that is not yet contained in tximeta’s built-in hash table of reference transcriptomes. For such cases, we have developed functionality in tximeta to formally link a given hashed checksum to a publicly available FASTA file(s) and a GTF/GFF file. The `makeLinkedTxome` function can be called, pointing to the transcriptome index as well as to the locations of the FASTA files and GTF/GFF file, and this will perform two operations: (1) it will add a row to a custom hash table, managed by BiocFileCache, and (2) it will produce a JSON file that can be shared or uploaded to public repositories, which links the transcriptome checksum with the source locations. When the JSON file is provided to `loadLinkedTxome` on another machine, it will add the relevant row to tximeta’s custom hash table, so tximeta will then recognize and automatically populate metadata in a similar manner to if the checksum matched with a transcriptome in tximeta’s built-in hash table. Finally, the cache location for tximeta, managed by BiocFileCache, can be shared across users on a cluster, for example, such that parsed databases, range objects, and custom hash tables created by any one user can be leveraged by all other users in the same group.

## Results

### Importing quantification data from known transcriptome

An example of importing RNA-seq quantification data using tximeta can be followed in the tximeta or fishpond Bioconductor package vignettes. Here we demonstrate the case where the Salmon files were quantified against a transcriptome that is in tximeta’s pre-computed hash table (for a list of supported transcriptomes as of the writing of this manuscript, see Table 1). Import begins by specifying a sample table (the “column data”, as the columns of the SummarizedExperiment object correspond to samples from the experiment).

~~~
coldata <-read.csv(“coldata.csv”)
~~~

For example, in the fishpond Bioconductor package vignette [32], the following `coldata` is read into R in the beginning of the analysis (here just showing the first two rows and five columns). The samples are from a human macrophage RNA-seq experiment [33].

~~~
##           names sample_id line_id replicate condition_name
## 1 SAMEA103885102 diku_A diku_1 1 naive
## 2 SAMEA103885347 diku_B diku_1 1 IFNg
~~~

This table must have a column `files` which points to paths of quantification files (`quant.sf`), and a column `names` with the sample identifiers. The following line can be used to create the `files` column (if it does not already exist), where `dir` specifies the directory where the Salmon output directories are located, and here assuming that the sample names have been used as the Salmon output directory names.

~~~
coldata$files <-file.path(dir, coldata$names, “quant.sf”)
~~~

It is expected that the quantification files are located within the original directory structure created by Salmon and with all the associated metadata files. The next step is to provide this table to the `tximeta` function, which returns a SummarizedExperiment object.

~~~
se <-tximeta(coldata)
~~~

Tximeta will print the following messages as the files are being imported (if a match is found, and if this checksum has been seen before by tximeta).

~~~
## importing quantifications reading in files with read_tsv ## found matching transcriptome:
## [ GENCODE - Homo sapiens - release 97 ]
## loading existing EnsDb created: 2019-01-01 12:34:56
## loading existing transcript ranges created: 2019-01-01 12:34:56
~~~

If the checksum matched one of the custom transcriptome checksums that was created or loaded by the user, the function would report, “`found matching linked transcriptome`”. A demonstration of such a workflow is given in the following section.

The SummarizedExperiment object, `se`, that is returned by tximeta can then be passed to various downstream packages such as DESeq2, edgeR, limma-voom, or fishpond, with example code in the tximeta package vignette [32, 34–37]. The transcript or gene ranges can be easily manipulated using the GenonicRanges or plyranges packages in the Bioconductor ecosystem [29, 38]. For example, to subset the object to only those transcripts that overlap a range defined in a variable `x`, the following line of code can be used.

~~~
se_sub <-se[se %over% x,]
~~~

The metadata columns associated with the genomic ranges of the SummarizedExperiment will have different information depending on the source. For GENCODE, Ensembl, and RefSeq, the chromosome names, start and end positions, strand, and transcript or gene ID are always included. Quantification data with an Ensembl source will also include the transcript biotype, and the start and end of the CDS sequence in the metadata columns.

Further examples of manipulating the SummarizedExperiment object can be found in the tximeta vignette, in the fishpond vignette, and in the plyrangesTximetaCaseStudy package [39].

### Importing data from a *de novo* transcriptome

It is also possible to use tximeta to import quantification data when the transcriptome does not belong to those in the set covered by pre-computed checksums (Table 1). This case may occur because the reference transcriptome is from another source or another organism than those currently in this pre-computed set, or because the transcriptome has been modified by the addition of non-reference transcripts (e.g. cancer fusion transcripts, or pathogen transcripts) which changes the checksum, or because the entire transcriptome has been assembled *de novo*. In all of these cases, tximeta provides a mechanism for local metadata linkage, as well as a formal mechanism for sharing the link between the quantification data and publicly available reference transcriptome files.

The key concept used in the case when the checksum is not part of the pre-computed set, is that of a link constructed between the transcriptome used for quantification via its hashed checksum and publicly available metadata locations (i.e. permalinks for the FASTA and GTF/GFF files). This link is created by the tximeta function `makeLinkedTxome` which stores the reference transcriptome’s checksum in a custom hash table managed by BiocFileCache, along with the permalinks to publicly available FASTA and GTF/GFF files.

We demonstrate this use case with an RNA-seq experiment [40] of transcripts extracted from the speckled killifish (*Fundulus rathbuni*) quantified using Salmon [24] against a *de novo* transcriptome assembled with Trinity [41] and annotated via dammit [42]. An example workflow is provided in the denovo-tximeta repository on GitHub [43]. Here, the FASTA sequence of the *de novo* assembly as well as a GFF3 annotation file have been posted to Zenodo [44, 45], and permalinks are used to point to those records. After the reference transcripts have been indexed by Salmon, the following tximeta function can be called within R.

~~~
makeLinkedTxome(
   indexDir=“F_rathbuni.trinity_out”,
   source=“dammit”,
   organism=“Fundulus rathbuni”,
   release=“0”,
   genome=“none”,
   fasta=“https://zenodo.org/record/1486276/files/
            F_rathbuni.trinity_out.fasta”,
   gtf=“https://zenodo.org/record/2226742/files/
            F_rathbuni.trinity_out.Trinity.fasta.dammit.gff3”,
   jsonFile=“F_rathbuni.json”
)
~~~

The function does not return an R object, but has the side effect of storing an entry in the custom hash table managed by BiocFileCache, and producing a JSON file which can be shared with other analysts. The JSON file can be loaded with `loadLinkedTxome`, and it will likewise store an entry in the custom hash table of the machine where it is loaded. In either case, when the quantification data [40] is later imported using tximeta, the checksum will be recognized and the relevant metadata attached to the SummarizedExperiment object output.

~~~
coldata <-data.frame(files, names)
se <-tximeta(coldata)
## importing quantifications reading in files with read_tsv
## found matching linked transcriptome:
## [ dammit - Fundulus rathbuni - release 0 ]
## loading existing TxDb created: 2018-12-13 18:26:20
## generating transcript ranges
~~~

After running `tximeta`, the SummarizedExperiment object `se` will have attached to its rows the ranges described by the GTF/GFF object, including any metadata about those transcripts. In the case of the killifish RNA-seq experiment, the transcript ranges have length, strand, and an informative column `gene id`. The ranges of the SummarizedExperiment can be examined (here only showing the first two ranges, and suppressing range names).

~~~
rowRanges(se)
## GRanges object with 143492 ranges and 3 metadata columns:
##         seqnames ranges strand | tx_id
##            <Rle> <IRanges> <Rle> | <integer>
##   TRINITY_DN114791_c0_g1_i1   1-2308   + | 1290
##   TRINITY_DN114724_c0_g2_i1   1-635    - | 1283
##                                gene_id
##                         <CharacterList>
##   ORF Transcript_…type:complete len:190 (+)
##   ORF Transcript_…5prime_partial len:83 (-)
~~~

## Availability and Future Directions

We outline an implementation for importing RNA-seq quantification data that involves (1) the quantification tool (here, Salmon) computing a hashed checksum of the reference transcript sequences, which are embedded in the index and in the per-sample output metadata, followed by (2) downstream comparison of checksums with a hash table (here, by tximeta), automated downloading and parsing of the appropriate metadata, and attachment to a rich object that bundles data and reference sequence metadata. The software is implemented within the R/Bioconductor environment for genomic data analysis, and leverages a number of existing Bioconductor packages for parsing annotation files, metadata storage, and genomic range manipulation [27, 29–31]. The tximeta package is available at https://bioconductor.org/packages/tximeta.

Currently, the pre-computed hashed checksums are focused on human, mouse, and fruit fly reference transcripts, from the popular reference transcriptome sources GENCODE, Ensembl, and RefSeq. Additional transcriptome releases from these sources are programmatically downloaded, the hashed checksum computed, and the checksum added to the tximeta package on Bioconductor’s 6 month release cycle. We are hopeful that future integration of tximeta with reference sequence retrieval efforts from the GA4GH consoritium will allow for a wide expansion of the number of supported organisms. Potentially all of the releases of reference transcriptomes from Ensembl and/or RefSeq may be supported by a future reference sequence retrieval API (GENCODE releases since 2015 are already fully supported by tximeta). Furthermore, we provide a mechanism for formally linking those reference transcripts not in any pre-computed hash table (e.g. *de novo* transcriptomes) with publicly available metadata. Finally, we plan to develop tximeta to support provenance identification at the level of alleles, by combining our current reference transcript identification with transcript variant identification as described in GA4GH’s Variant Representation Specification [11].

All bioinformatic software packages have a finite lifespan, including the package described here. We join with others in recommending the underlying paradigm of embedding reference sequence checksums in sample output metadata, followed by downstream database lookup of checksums, and identification of reference sequence metadata. This paradigm should be adopted by other bioinformatic software that outputs any data that refers to a reference sequence. This workflow has the advantage of not requiring additional effort or actions on the part of the upstream bioinformatic analyst. Otherwise, we risk exposing downstream analysts to the “major time thief” of *post hoc* guesswork involved in identifying the provenance of datasets shared publicly but without critical metadata [7].

## Data Availability

All datasets used in this manuscript are available as Bioconductor data packages used in the tximeta or fishpond package vignettes, or in the case of the *de novo* transcriptome analysis, have been deposited to Zenodo:

- Quantification data (Salmon output directory, tar.gz) [40] DOI: 10.5281/zenodo.1486283
- De novo transcriptome assembly (FASTA) [44] DOI: 10.5281/zenodo.1486276,
- Annotation file (GFF3) [45] DOI: 10.5281/zenodo.2226742,

## Acknowledgments

The authors thank the following individuals for useful discussions in the development of tximeta: Vince Carey, Paul Flicek, Joel Parker, Oliver Hoffmann, Stephen Turner, Shannan Ho Sui, Thomas Keane, Andy Yates, Reece Hart, Matthew Laird, Terence Murphy, Nathan Sheffield. The authors also thank Reid Brennan, and C. Titus Brown, and Andrew Whitehead for allowing use of the killifish transcriptome dataset in the *de novo* transcriptome example.

